# Activity of the GHRH Antagonist MIA602 and its Underlying Mechanisms of Action in sarcoidosis

**DOI:** 10.1101/2020.08.19.257915

**Authors:** Chongxu Zhang, Runxia Tian, Emilee M Dreifus, Gregory Holt, Renzhi Cai, Anthony Griswold, Pablo Bejarano, Robert Jackson, Andrew V. Schally, Mehdi Mirsaeidi

## Abstract

Growth hormone releasing hormone (GHRH) is a potent stimulator of GH secretion from the pituitary gland. Although GHRH is essential for the growth of immune cells, the regulatory effects of its antagonist in granulomatous disease remains unknown. Here, we report expression of GHRH receptor (R) in human tissue with sarcoidosis granuloma and demonstrate the anti-inflammatory effects of MIA602 (a GHRH antagonist) in two *in vitro* human granuloma models and an *in vivo* granuloma model. MIA602 decreases levels of IL2, IL12, and IL17A in *in vitro* granuloma model.

We show further that the anti-inflammatory effect of MIA602 appears to be mediated by reduction in CD45^+^+CD68^+^ cells in granulomatous tissue and upregulation in PD-1 expression in macrophages.

In analysis of expression of proteins involved in the mitochondrial stage of apoptosis, we show that MIA602 increases the levels of caspase 3, BCL-xL/BAK dimer, and MCl-1/Bak dimer in granuloma. These findings indicate that MIA602 may not induce apoptosis.

The clinical relevance of our findings further suggest that HGRH-R is potentially a target for treatment of granulomatous disease and MIA602 possibly a novel therapeutic agent for sarcoidosis.

## Introduction

Sarcoidosis is an inflammatory disease characterized pathologically by noncaseating granulomas (1). Sarcoidosis affects every race and ethnicity, but in the U.S., it is more commonly seen in African Americans (2). Sarcoidosis is triggered by environmental factors such as dust, mold, mycobacteria, and occupational exposures, in addition to a genetic predisposition (3, 4).

No known medications specifically treat sarcoidosis, but corticosteroids such as prednisone and prednisolone have been used due to their anti-inflammatory properties. Despite corticosteroids showing some positive treatment results, many patients continue to experience significant problems because of the development of fibrosis from previously active or smoldering granulomatous inflammation (5, 6). In addition, many patients report significant steroid-associated side effects (7). Therefore, there is a great need for sarcoid-specific medications to target inflammatory pathways and prevent sarcoidosis sequelae. If inflammation can be modulated, this would likely cause a decrease in symptoms associated with sarcoidosis, leading to disease remission.

Growth-hormone releasing hormone (GHRH) is a hypothalamic 40-44 amino acid peptide hormone that specifically stimulates release and synthesis of growth hormone (GH) from the pituitary gland (8). Growth hormones are active throughout the body and promote growth and development. The GHRH receptor (GHRH-R) is a seven transmembrane subunit G-protein-linked receptor found predominantly in the pituitary gland (9). HRH-R can be blocked by synthetic peptides known as GHRH-R antagonists. These peptides include the MIA and AVR classes of antagonists (10). MIA602 has been shown to modulate lung inflammation and fibrosis (11). GHRH-R antagonists function in a cAMP-dependent pathway and GHRH-R antagonists downregulate p21 activated kinase 1 (PAK1)-mediated signal transducer and activation of transcription factor 3 (STAT3)/nuclear factor kB (NFkB) (12). To date, however, none of GHRH-R antagonists have known anti-inflammatory properties in granulomatous diseases or sarcoidosis (13).

In this study, we investigated the anti-inflammatory properties of MIA602 by measuring cytokines and gene expression in both an *in vitro* and an *in vivo* granuloma models. We developed the *in vitro* model by exposing human peripheral blood mononuclear cells (PBMCs) to microparticles generated from mycobacterial cell walls (14). We expose mouse lungs to the same microparticles to study lung granulomatous disease in vivo (15).

## Results

### MIA602 has anti-inflammatory effects in an *in vitro* sarcoid-like granuloma model

To determine if MIA602 can reduce inflammation in granulomatous disease, we first asked if GHRH-R is expressed in sarcoidosis granuloma. To address this question, we first stained 6 human lung samples with pathological confirmation of sarcoidosis for GHRH-R using Immunohistochemistry (IHC). As shown in Figure 1, granulomata significantly expressed levels of GHRH-R.

**Figure 1.**
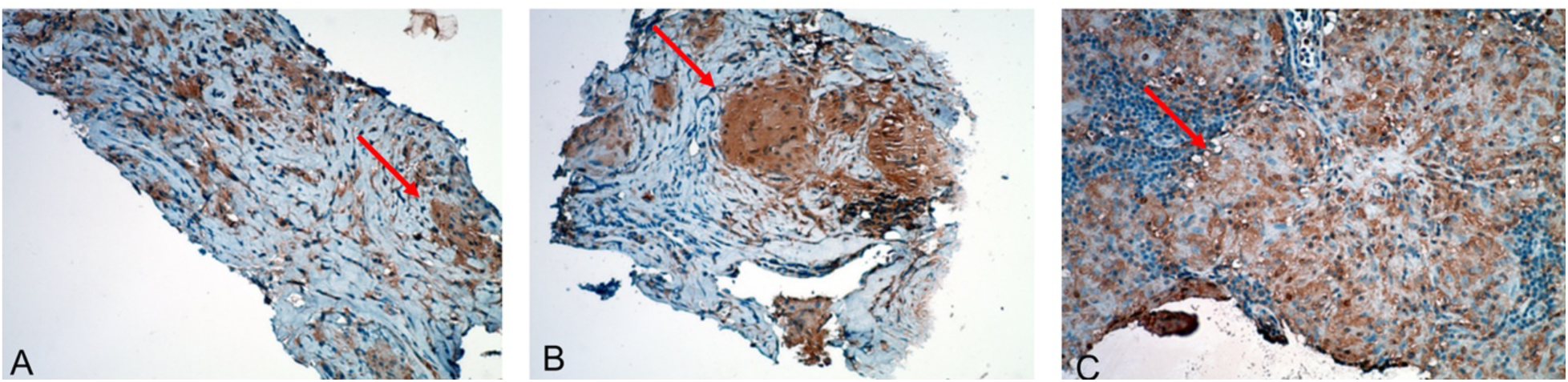
Lung histopathology. Lymph node tissue from 3 subjects with confirmed sarcoidosis were recut. 5 µm sections were stained for GHRHR. The arrow shows granuloma. The Brown color in the tissue indicates expression of GHRHR.

After confirming the presence of GHRH-R in granuloma, we aimed to determine anti-inflammatory properties of the GHRH-R antagonist (MIA602) in an *in vitro* sarcoid-like granuloma system. To achieve this goal, we used PBMC from 5 confirmed subjects with sarcoidosis who participated in an IRB-approved sarcoidosis biobank. Samples of PBMC were challenged with microparticles generated from *M. abscessus* cell wall (called microparticles in this manuscript) and treated with saline, 1 μM MIA602, or high dose methyl prednisolone (138 μM) and compared to control PBMCs. PBMCs challenged with microparticles all formed granulomas. Medium was removed 48h after treatments and cytokines were measured using a multiplex Elisa instrument. We found a significant difference in expression of several cytokines in the granuloma group in comparison with control. MIA602 significantly reduced production of IL2, IL12, and IL17A, as shown in Figure 2. In comparison with steroids, MIA602 showed a significant anti-inflammatory effect.

**Figure 2.**
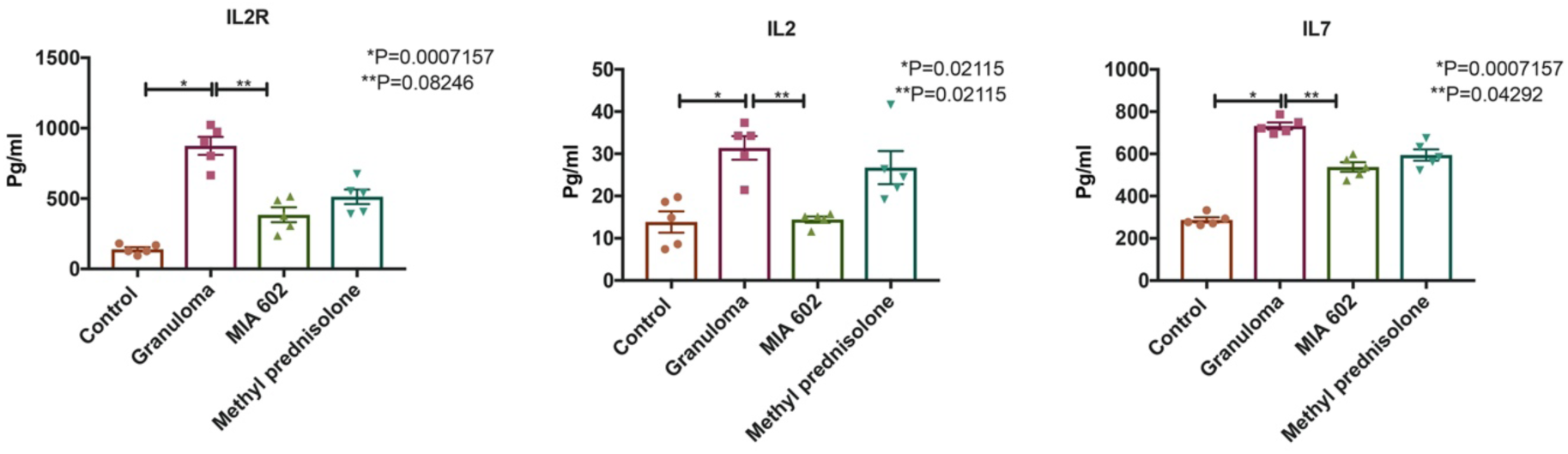
Shows ELISA results for in vitro sarcoid-link granuloma. Granuloma was developed using PBMC from subjects with confirmed sarcoidosis. Control: PBMC, Granuloma: PBMC challenged with microparticles, MIA602: Granuloma treated with MIA602, Methyl prednisolone and Granuloma treated with methyl prednisolone. Data shown are mean cytokine concentrations ± SEM at 48 hours after challenge with MAM microparticles (N=5)

### MIA602 does not induce apoptosis in an *in vitro* sarcoid-like granuloma model

We tested the hypothesis that MIA602 may influence respiration and apoptosis in granuloma cells. The specific processes of apoptosis and its relation to mitochondrial dynamics in granuloma have not been fully elucidated. To test if MIA602 has apoptotic effects, we measured the protein levels of pro- (active caspase-3) and anti-apoptotic factors (survivin, Bcl-xL/Bak dimer, and Mcl-1/Bak dimer) in the *in vitro* granuloma model. PBMC samples from 5 confirmed sarcoidosis subjects were challenged with microparticles and treated with saline, 1 μM MIA602, methyl prednisolone (138 μM), and control PBMCs. Medium samples were removed 48h after treatments.

As shown in Figure 3, the level of Mcl-1/Bak dimer was significantly decreased in granuloma but was restored by MIA602. BclxL/Bak Dimer levels increased in granuloma treated with MIA602 in comparison with granuloma treated with saline. Active caspase 3 level also increased significantly in granuloma compared to PBMC possibly due to lymphocyte early activation. MIA602 further increased levels of active caspase 3. Survivin levels showed no difference between groups.

**Figure 3.**
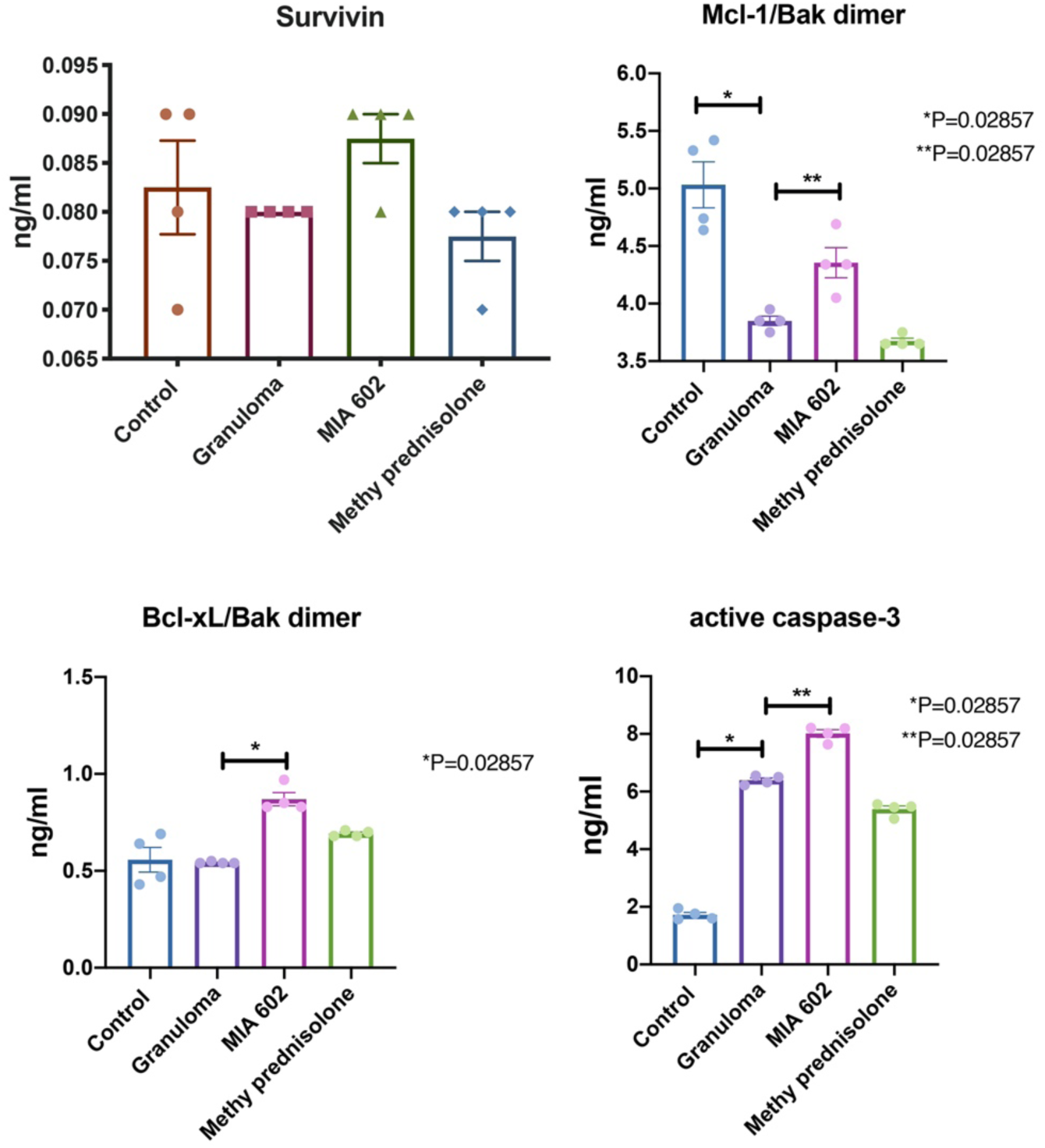
Shows ELISA the protein levels of pro- (active caspase-3) and anti-apoptotic factors (survivin, Bcl-xL/Bak dimer, and Mcl-1/Bak dimer) in the *in vitro* granuloma model. Granuloma developed from PBMC samples of 5 subjects with sarcoidosis after challenging with microparticles generated from *M. abscessus* cell wall. Control group is PMBC samples from the same subjects that were not challenged. A granuloma group was treated with 1 μM MIA602, and another with methyl prednisolone (138 μM). The PBMCs were collected and lysed and protein was extracted from each group separately for ELISA (n=5).

### MIA602 reduces inflammation in the lung of a sarcoidosis mouse model

We established a mouse pulmonary granulomata model from exposure to the microparticles and have used it to explore the role of type I IFN pathways in inflammation. This model is applicable to pulmonary sarcoidosis studies due to the similarly of granuloma to human sarcoidosis. We used C57Bl/6 mice to develop the model as presented in detail elsewhere (15). Granulomatous disease was induced by addition of the microparticles in the lower respiratory tract of mice, and we confirmed that the expression of GHRH-R was significantly increased in the lungs with granulomata. Mice were treated with subcutaneous injection of 5 µg MIA602 daily. After 3 weeks, mice were sacrificed and inflammation graded by a lung pathologist. Lung samples were stained with H&E, CD68, PD1, PD-L1, and CD30 and scored based on the percentage of cells expressing each marker. As shown in Figure 4, the granuloma group had significant inflammation in the lung, but the mice treated with MIA602 had lower inflammation scores. Figure 4B shows immunofluorescence staining of MIA602 in the granulomatous reaction of mice lung.

**Figure 4.**
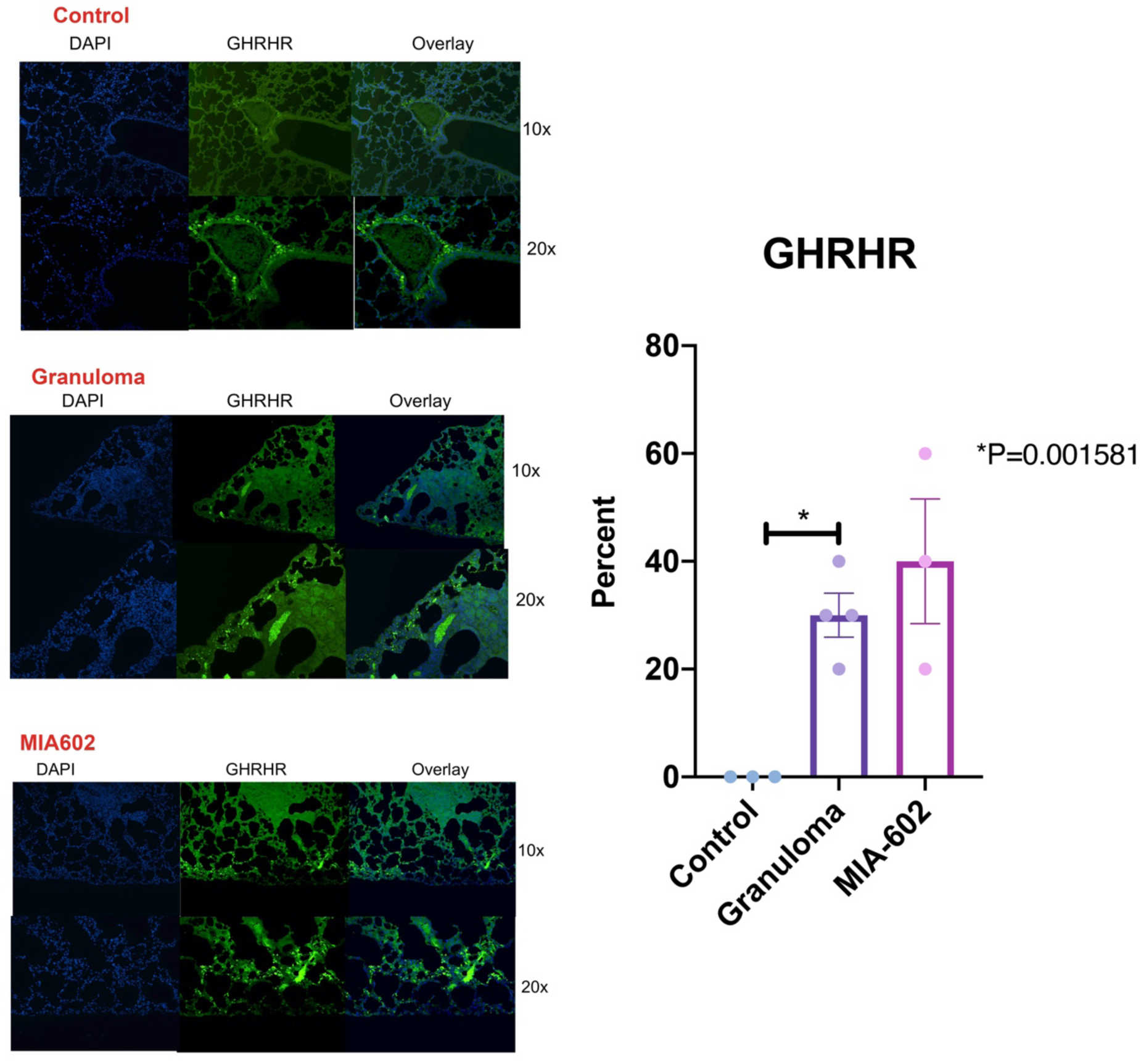
A representative image immunofluorescence microscopic detection of GHRHR in lungs of mice Magnification are 10X and x20 for all representative images. *P*-value shows percentage differences of lung stained cells between challenged mice and controls (N= 3 controls, 4 challenged, and 3 MIA602 mice). Control mouse was not challenged, granuloma group challenged with microparticles generated from MAB cell wall, MIA602 mice were challenged with granuloma and received 5 μg MIA602 daily injection. The staining intensity and percentage of area extent of the immunohistochemistry was evaluated by a lung pathologist. A: control B: challenged with microparticles with developed sarcoid-like granuloma.

The inflammation score of lungs showed a higher percent inflammation in the granuloma group treated with saline. Lung inflammation was almost normalized in mice that received treatment with MIA602 as shown in **Figure 5**. CD30^+^ cells were statistically significantly increased in the lungs of mice treated with MIA602, which also showed reduced number and size of granuloma. The percentage of CD68^+^ cells in the lung increased in granuloma and decreased in MIA602 (not statistically significant). This same pattern can also be seen in PD-1^+^- and PD-L1^+^ cells in granuloma (treated with saline) and MIA602 groups.

**Figure 5.**
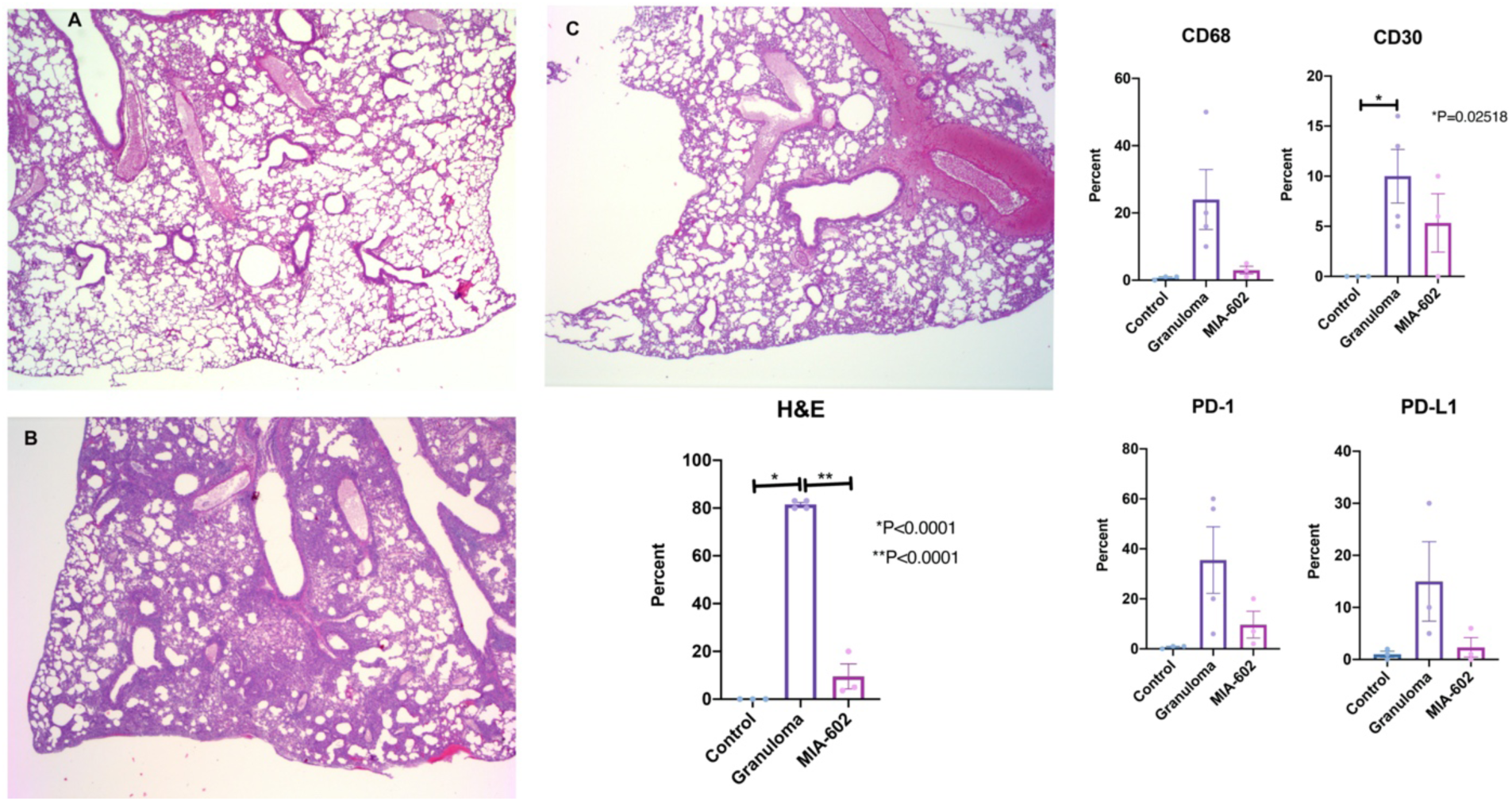
Shows representative images for H&E and IHC changes in the lungs *P*-value shows percentage differences of lung stained cells between challenged mice and controls (N= 3 controls, 4 challenged, and 3 MIA602 mice). Control mouse was not challenged, granuloma group challenged with microparticles generated from MAB cell wall, MIA602 mice were challenged with granuloma and received 5 μg MIA602 daily subcutaneous injection. The percentage of area extent of the immunohistochemistry was evaluated by a lung pathologist. Representative images for H&E are shown in A (control) B (granuloma) C (treatment with MIA602). IHC images were not shown.

Mice were challenged with the microparticles and treated with saline, MIA602 (5 μg), or methyl prednisolone (100 μg) and were compared to control mice. After 3 weeks, lungs were harvested from all groups and single cells were generated for flow cytometry analysis. As shown in Figure 6, the population of CD45+CD68 cells significantly increased in challenged mice. MIA602 treatment significantly reduced these cell population. As it has been previously shown that PD1 and PD-L1 play important roles in granuloma formation (16), we hypothesized that MIA602 would restore the number of CD68+ cells that express PD-1 and PD-L1. Figure 6 shows that CD45+CD68+ cells that expressed PD-1 were indeed significantly reduced in mice challenged with granulomatous reaction. The percentage of cells significantly increased with MIA602 and reached to the levels of the control group.

**Figure 6.**
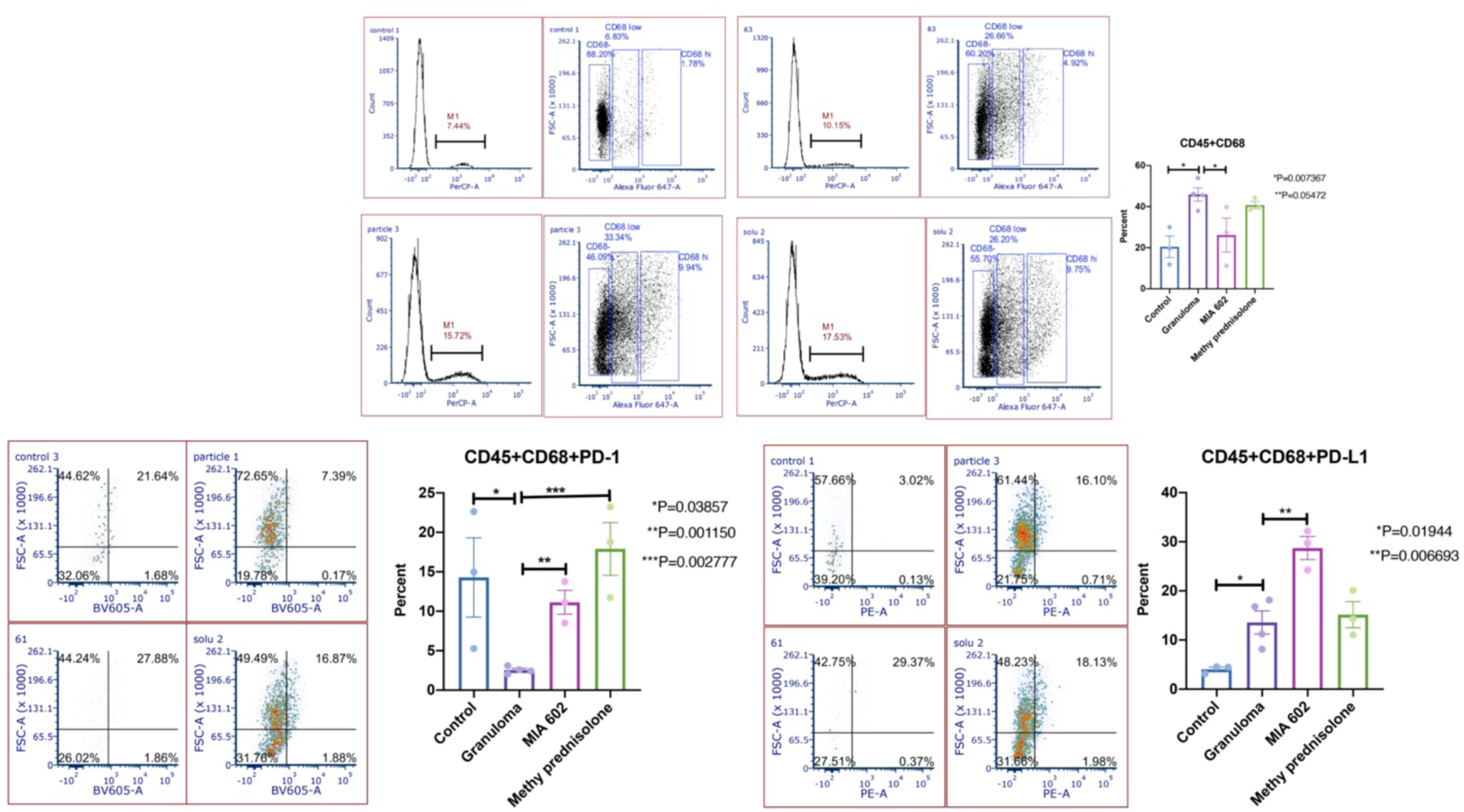
Flow cytometry analysis of single cells generated from mice lung. Gated the nucleated cells for the size and complexity of the event then plotted in FSC-A and FSC-H to gate single cells and exclude doublets. The live leukocytes were defined as CD45^+^ and the CD45^+^CD68^+^ population was defined as macrophage. A. control, B. mice challenged with microparticles generated from MAB cell wall to develop granuloma, and C. mice challenged with microparticles that treated with daily subcutaneous injection of 5 µg MIA602. (N= 3 controls, 3 challenged, and 3 MIA602 mice).

To confirm that mouse lungs with granuloma showed a higher percentage of CD45+CD68+ cells that expressed PD-L1, we stained lung single cells in all experimental groups. As shown in Figure 6, the population of CD45+CD68+PD-L1 cells was significantly increased in granuloma and further increased after treatment with MIA602. Thus, we conclude that the anti-inflammatory effects of MIA602 extend mainly through reduction in CD68+ cells and upregulation of PD-1 expression.

### Inducible Nitric Oxide Synthase and Nitrotyrosine are increased in lungs of mice with sarcoidosis

Inducible nitric oxide synthase (iNOS) produces nitric oxide and plays a role in granuloma development as macrophages increase expression of iNOS after an exposure to bacterial lipopolysaccharides and IFNγ (17). We have previously shown that GHRH also increases the expression of the iNOS (18). While the effects of GHRH-R antagonists in oxidative stress have been explored previously (19), the effects of these peptides in nitrosative stress have not been well evaluated. To understand effects of MIA602 on the nitric oxide response, we measured iNOS and nitrotyrosine (as an indicator of NO function), in mouse lungs that were challenged with the microparticles (sarcoidosis model) and injected daily with saline, daily subcutaneous injections of 5 μg MIA602, and we compared them to the control group who were never challenged. Lung tissue was harvested after 3 weeks and stained with NOS2 and nitrotyrosine. As shown in Figure 7, expression of NOS2 was increased in sarcoid-like granulomas and statistically unchanged in MIA602 treated mice.

**Figure 7.**
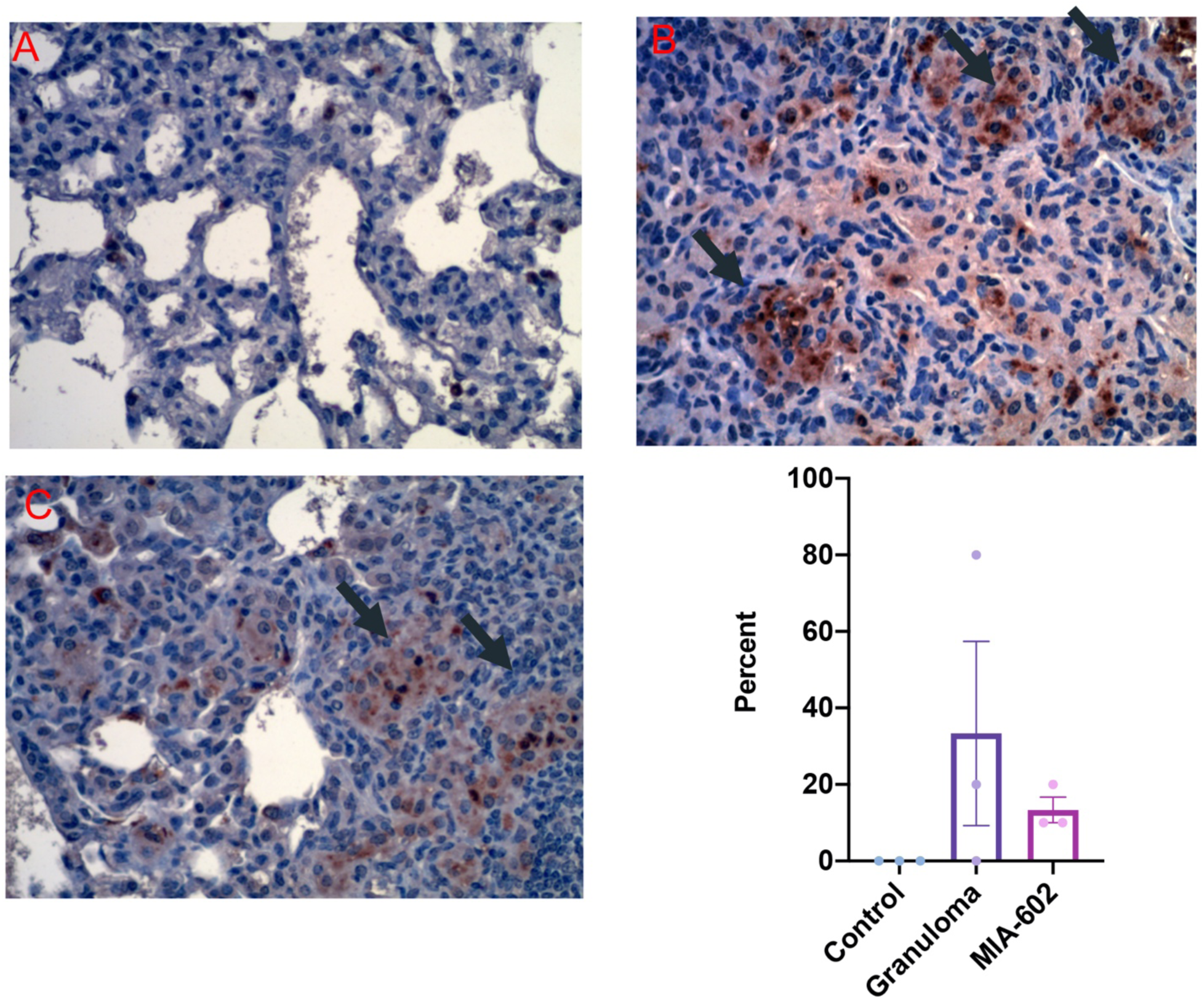
Shows A representative image of Immunohistochemical staining for iNOS2 staining in the lung of mice. A. control, B. mice challenged with microparticles generated from MAB cell wall to develop granuloma, and C. mice challenged with microparticles that treated with daily injection of 5 µg MIA602. The black arrow shows granuloma with iNOS2 staining. (N= 3 controls, 3 challenged, and 3 MIA602 mice). The Brown color in the tissue indicates expression of iNOS2. P>0.05 (not statistically significant).

Nitrotyrosine expression was statistically significantly increased in the lungs of mice with sarcoid-like granuloma as shown in Figure 8. Treatment with MIA602 decreased the nitrotyrosine levels but was not significant statistically. This confirms that mouse lung after challenge with microparticles activates iNOS and increases nitrotyrosine. MIA602 reduced both, but not statistically significantly, in our experiments. This experiment demonstrates a mild anti-nitrosative effect of MIA602 in the sarcoidosis mouse model.

**Figure 8.**
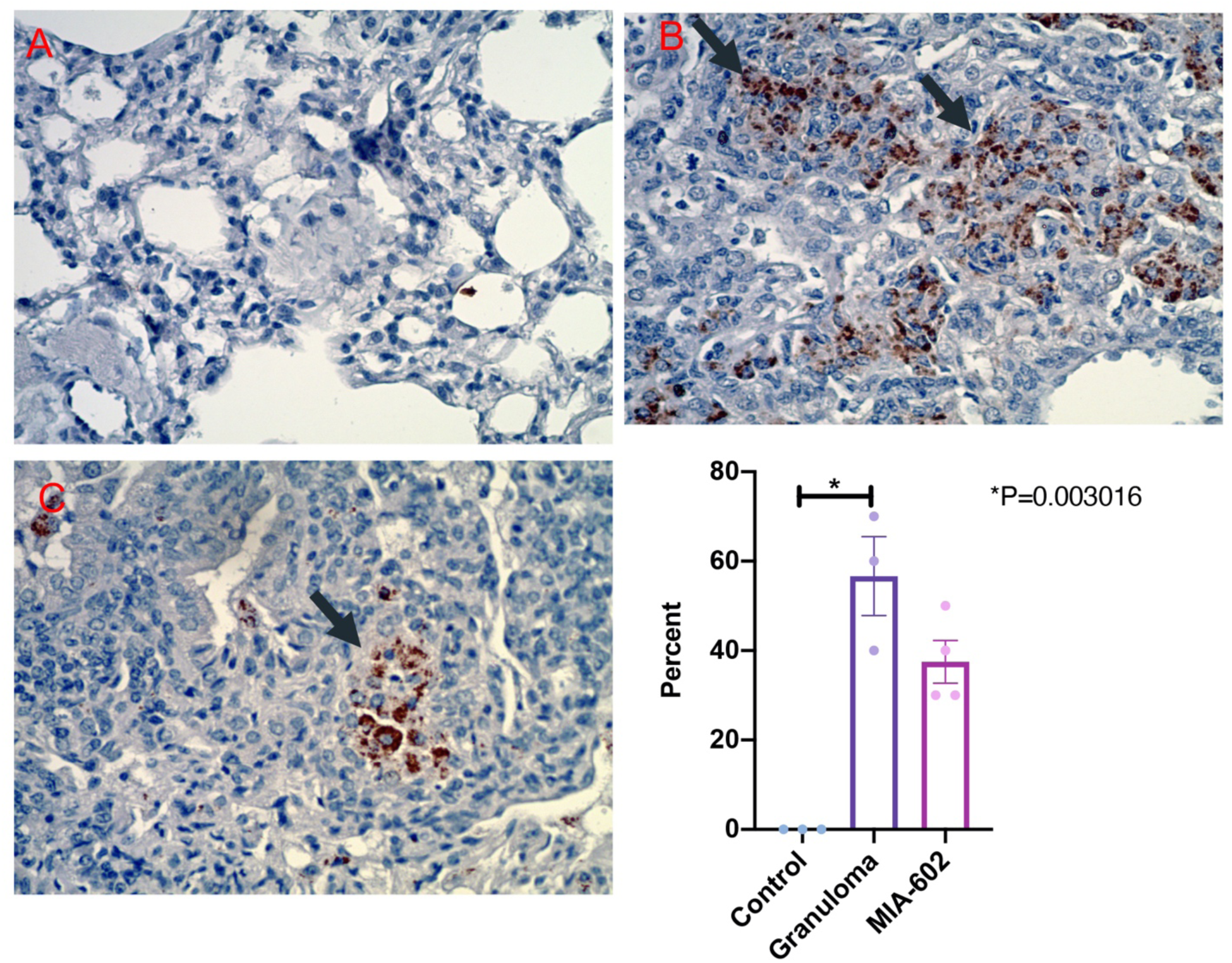
Shows representative Immunohistochemical staining for nitrotyrosine in lung of mice A. control, B. mice challenged with microparticles generated from MAB cell wall to develop granuloma, and C. mice challenged with microparticles that treated with daily injection of 5 µg MIA602. The black arrow shows granuloma with nitrotyrosine staining. (N= 3 controls, 3 challenged, and 3 MIA602 mice). The Brown color in the tissue indicates expression of nitrotyrosine.

### MIA602 significantly changed transcriptomics in sarcoidosis mouse model

We extracted RNA from lungs challenged with the microparticles and treated with saline daily or 5 μg MIA602. RNA was isolated from lung tissue 3 weeks after challenge and RNASeq was performed.

Overall, we identified 1407 protein coding genes differentially expressed between Granuloma and MIA602 with a false discovery rate p-value < 0.05 (599 upregulated in MIA602, 808 downregulated in MIA602). In order to identify the most robustly changing genes, we extracted only those with at least a 2-fold change and identified 489 genes (226 upregulated in MIA602, 263 downregulated in MIA602). Genes with a twofold change are visualized in a heatmap (Figure 9) showing consistent and robust differences between these two groups of samples.

**Figure 9.**
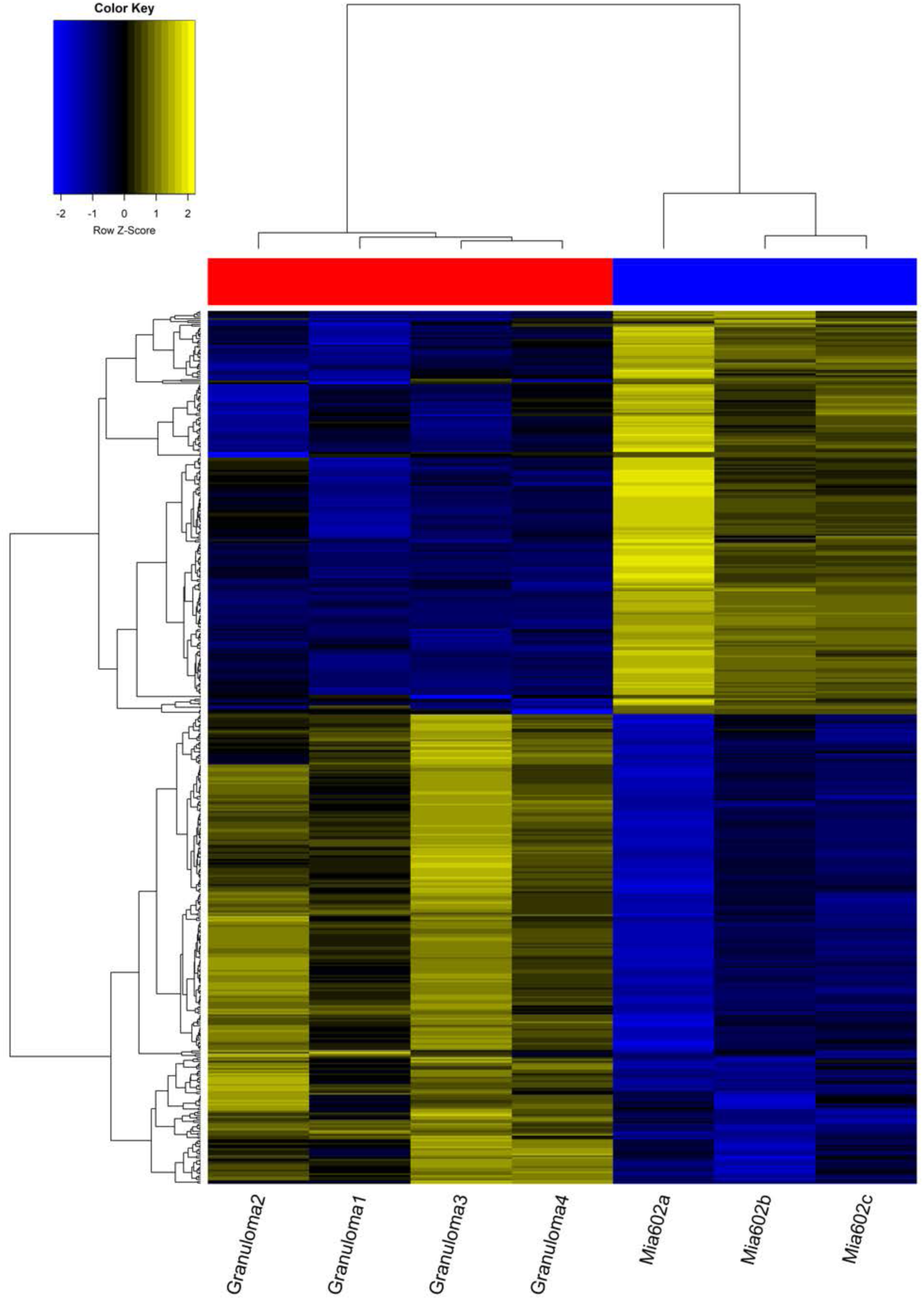
shows heatmap of genes with FC>2.5. Coding genes was illustrated in the heatmap between mice lung with granuloma treated with saline vs. mice lung with granuloma treated with MIA602. P: challenged with the microparticles (with pulmonary granulomata) and injected daily with saline, MIA602: challenged with microparticles and treated with daily subcutaneous injection of 5 µg MIA602. Lung of mice with granulomatous inflammatory reaction treated with MIA602 in comparison with saline differentially expressed several coding genes. (N=3 mice treated with MIA602, N=4 mice challenged and treated with saline)

## Discussion

MIA602 is a novel, potentially therapeutic agent for sarcoidosis. We found that MIA602 has an anti-inflammatory effect in an in vitro sarcoidosis model with significant reduction in inflammatory cells. We show further that the effect of MIA602 may not be due to apoptosis in granulomata. The anti-inflammatory effect of MIA602 appears to be mediated by a reduction in CD45^+^+CD68^+^ cells in granulomatous tissue and upregulation in PD-1 expression in macrophages. We also showed significant changes in gene profiles of mouse lungs treated with MIA602 vs. saline.

The anti-inflammatory effects of MIA602 in granuloma is a novel finding consistent with previous reports on anti-inflammatory properties of GHRH-R antagonists in pulmonary fibrosis (11), ocular inflammation (20), and chronic prostatitis (21).

The current study suggests that MIA602 may alleviate inflammatory response by reduction of central cytokines including IL-17A in sarcoidosis. IL2, IL12, and IL17 are major cytokines in the pathogenesis of sarcoidosis (22). Prasse *et al* demonstrated that bronchoalveolar lavage fluid of patients with sarcoidosis has high concentrations of IL-2 and IFN-γ (23). The roles of TH17 cells and IL17A in sarcoidosis are well known (24). The in vivo experiments in the sarcoidosis mouse model show that tissue inflammation scores were significantly reduced by MIA602 treatment, and this effect is mainly macrophage dependent. GH has an important role in maturation and activation of macrophages (25). While the role of GHRH agonist on T cells was described previously by Khorram *et al (26)*, the current study suggests an anti-inflammatory role of the GHRH-R antagonist in a macrophage phenotype. Our IHC study shows that PD-1 expression in lung macrophages was significantly reduced in the granuloma sarcoid model, and MIA602 restored PD-1 expression. Thus, GHRH-R antagonist’s anti-inflammatory effects may be partially due to expression of PD-1+ macrophages. Gordon *et al*. suggested that PD-1+ macrophages may be associated with type 2 polarization and suppression of phagocytosis (27). The M2 phenotype of macrophages is known to have an immunosuppressive effect with anti-inflammatory cytokine secretion to allow wound healing (28). PD-L1+ macrophages increased in the lungs of mice with sarcoidosis like granulomas, and treatment with MIA602 induced higher numbers of PD-L1 macrophages. The functionality of PD-L1 in macrophages is not well understood. It has been suggested that macrophages are destroyed by T cells via PD-L1 (29). Our study has several limitations. The effect of MIA602 in neutrophils and lymphocytes has not been evaluated in our study. Lymphocytes are important effectors in granuloma. The role of a ratio between neutrophils and lymphocytes in sarcoidosis has been discussed elsewhere (30). The effect of MIA602 in caspase 3 needs further investigation. MIA602 significantly increased active caspase 3 concentration. It may be associated with inducing apoptosis in granuloma, or may be due to early activation of stimulated T cells (31). MIA602 reduced iNOS, but not statistically significantly, in our experiment. This finding could be due to limitations in our sample size.

The current study shows that MIA602 has important anti-inflammatory properties through reduction in IL2, IL12, and IL17A release and reducing CD68^+^ cells in the lung. These effects may be mediated through checkpoint receptors (PD1 and PD-L1). Further investigation will lead to understanding the anti-inflammatory effects of MIA602 in sarcoidosis and explore its potential therapeutic role.

## Methods

### MIA602 preparation

MIA602 was prepared by Dr. Renzi Cai and Dr. Andrew Schally. The chemical structure of MIA602 is [PhAc-Ada0, D-Arg2, Fpa56, Ala8, Har9, Tyr (Me)10, His11, Orn12, Abu15, His20, Orn21, Nle27, D-Arg28, Har29] hGH-RH (1–29) NH2. MIA602 was dissolved in 100% dimethyl sulfoxide (DMSO, ACS grade, Sigma) for stock and diluted at 1:1000 in corresponding culture medium to a final concentration of 1 μM before use as previously presented. Control group *in vitro* and *in vivo* received placebo with the same volume and concentration of DMSO.

### Microparticle development

As previously presented in details (14), microparticles were generated from a rough colony of a clinical strain of *mycobacterium abscessus* (MAB) with sonicating and heating live bacilli. High quality images of non-infectious, MAB particles were obtained by scanning electron microscope (SEM).

### Human blood sample

Blood samples were collected from 9 patients with confirmed pulmonary sarcoidosis and randomly selected from the University of Miami Sarcoidosis Biobanking, and matched by age, sex, and race with 10 healthy control by the University of Miami Institutional Review Board approval number 20150612. To avoid the inconvenience and risks associated with additional venipunctures, a 10 ml blood specimen was collected during an already scheduled venipuncture. Patients who currently had a hgb <7 mg/dL were excluded from participating in this study.

### Maturing *in-vitro* granuloma like formation

In vitro granuloma was developed from challenging PBMC to microparticles as previously described (14).

### Mouse Model Exposure to MAB Microparticles

Granulomatous reaction in the mouse lung was developed as previously described (15).

### ELISA

As we previously presented (14), PBMCs were lysed in lysis buffer (Cell Signaling Technology, Beverly, MA) with protease inhibitor cocktail (Cell Signaling Technology, Beverly, MA) and sonicated three times for 2 s each with at least 1-min rest on ice between each 2-s pulse. Samples were centrifuged at 10,000 × g for 5 min at 4°C and the supernatant was collected. Protein concentration was determined by BCA protein assay kit from Cell Signaling Technology. Thirty micrograms of total protein were mixed in a reducing sample buffer and used for mitochondrial apoptosis assay per the kit instruction. The assay was performed using a Bio-Rad kit, Cat# 171-WAR3CK.

To measure the cytokines in media, supernatant aliquot samples were analyzed, thawed, and spun at 12,000 rpm for 10 min to separate the particulate material at the bottom. 50 μl of undiluted media was plated from each sample onto a 96-well V-bottom plate (source plate) by manual pipetting according to predefined maps. The aliquots were wrapped in parafilm and kept in a humid chamber at 4 °C during the entire process, but not longer than 72 hr. Growth factors and their receptors’ capture antibodies were reconstituted and diluted per manufacturer specification and 50 μl plated into each well of respective 96-well high-binding half-well plates which were then sealed and incubated overnight at 4 °C. The cytokine levels were measured using a procartaplex human th1/th2 cytokine panel 11 plex from invitrogen, cat # epx110-10810-901.

#### Immunofluorescence confocal microscopy

*Detained methods were discussed elsewhere (15), As summary, mice were sacrificed on day 14, and the left lungs were harvested as presented earlier. Lungs were filled with 10% buffered formalin and fixed in formalin for at least 72 h before immunohistochemistry (IHC) staining. H&E staining was used to determine inflammatory pathology*.

For immunofluorescence, paraffin-embedded serial sections (5 μm) first underwent standard deparaffinization and rehydration procedures and were then probed with GHRHR (Origene, cat # TA311715) as primary antibody, and anti-Rabbit antibody from Sigma, (cat # F-9887) as secondary. Nuclei were counterstained with DAPI. All reagents were from Sigma-Aldrich. Tissue sections were analyzed using fluorescence microscopy and ImageJ software (version 6.0; NIH) to quantitate fluorescence intensity. In trichrome-stained slides, blue stain (collagen content) was also quantitatively analyzed using ImageJ.

Confocal immunofluorescence images were acquired using a Leica DM6000 microscope with a SP5 confocal module at the University of Miami McKnight Analytical Imaging Core Facility.Captured images were processed using Velocity Software version 6.1.1 software (Perkin-Elmer, Waltham, MA).

For immunohistochemistry,5-μm paraffin sections were processed by deparaffinization and rehydration followed by endogenous peroxidase blocking (1% H_2_O_2_ in methanol for 20 minutes) and antigen retrieval (boiled in 10 mM citrate buffer for 30 minutes). Tissue sections were blocked with 2% goat or horse serum (Vector Laboratories) and incubated with antibody CD68 (Proteintech, Cat# 25747-1-AP), PD-1 (cell signaling, Cat # 84651), PD-L1 (Proteintech, Cat# 17952-1-AP), CD30 (Lsbio, cat# LS-c162069) CD3 (cell signaling, cat# 99940), iNOS (invitrogen, cat #PAI-036), Nitrotyrosine (Novus, NBP2-54606) overnight at 4°C, washed with TBST five times, then secondary antibodies were added (Vector Laboratories, cat# PI-2000). Immunoreactivity was detected using the ABC Elite kit (Vector Laboratories). We used DAB as the final chromogen and hematoxylin as the nuclear counterstain. Negative controls for all antibodies were made by replacing the primary antibody with non-immunogen IgG. Lung inflammation was scored using the three fields with the highest infiltrate intensity at 100X power magnification as previously presented by our team (15). The area of inflammation was measured and averaged for the three examined high-power fields.

#### RNA isolation and analysis

RNA from mouse lungs was extracted using RNA Miniprep Plus Kit (Zymo Research). Briefly, the whole lung was homogenized in TRI reagent and total RNA extraction was performed following the instructions provided by the manufacturer with additional DNase treatment. Quantity and quality of the samples was determined by NanoDrop spectrophotometer and Agilent Bioanalyzer 2100, respectively.

Preparation and sequencing of RNA libraries was performed at the John P. Hussman Institute for Human Genomics Center for Genome Technology. Briefly, total RNA quantity and quality were determined using the Agilent Bioanalyzer. At least 300 ng of total RNA was used as input for the Nugen Universal Plus mRNA-Seq Library Preparation kit according to manufacturer’s protocol to create polyA enriched mRNA sequencing libraries. Sequencing was performed on the Illumina NovaSeq 6000, generating at least 40 million paired-end 100 base reads per sample.

Sequencing data were processed with a bioinformatics pipeline including quality control, alignment to the mm10 mouse reference genome, and gene quantification against the Ensembl build 91 mouse gene set. Count data was input into edgeR software for differential expression analysis. Counts were normalized using the trimmed mean of M-values (TMM) method to account for compositional difference between the libraries, and differential expression analysis between groups was performed using the quasi-likelihood F-test implemented in edgeR. Genes were considered to be statistically different with a false discovery rate p-value (FDR) ≤ 0.05. The raw RNAseq data is available as part of GEO SuperSeries GSE140439.

### Flow Cytometry

Mice were sacrificed on day 14 and the left lungs were harvested for pathology after perfusion of the right ventricle with 10 ml of PBS.

The upper half of left lung tissue (without trachea, main bronchus, or branches) was removed and rinsed by PBS to clean off blood. The tissue was minced and dispersed with scissors, to increase total surface area. To develop single cell suspension, we used the rubber end of a 5 ml plastic syringe to mesh cells through a 100 µm cell strainer with continuous rinse the cell strainer with ice-cold RPMI 1640 as shown in the Suppl Figure 1. Meshed cell suspension again through a 70 µm cell strainer and rinse thoroughly with 3 ml of washing buffer containing DNAse followed by 15 ml of DNAse-free washing buffer. Centrifuge at 286 x g and 18 °C for 5 min and discard the supernatant (32).

The cells (10^6^ cells/ml) were resuspended in 100μl protein blocking solution with 5 μl fluorescent conjugated antibodies, CD8 (Biolegend Cat# 100714, CD45 (Biolegend Cat# 103130), CD68 (Biolegend Cat# 137004), PD-1 (Biolegend Cat# 135219), PD-L1 (Biolegend Cat# 124308), CD4 (Biolegend Cat# 100510), CD11b (Biolegend Cat# 101243), CD11c (Biolegend Cat# 117318), F4/80 (Biolegend Cat# 123146), IFNg (Biolegend Cat# 505836). Samples were analyzed on a BD LSR II flow cytometer using BD FACSDiva software, and data analysis was performed using Flowjo software (TreeStar, Ashland, OR). Cell populations were identified using sequential gating strategy. The expression of activation markers is presented as median fluorescence intensity. Lung immune cells were classified based on FC marker expression as previously described (33).

### Statistical Analysis

A paired *t*-test was used to compare the values of means of two groups using GraphPad Prism 8 software. Data represent mean ± SEM, and results with *p* value (two-sided) less than 0.05 were defined as statistically significant.

### Study approval

Blood samples for this study were randomly selected from the University of Miami Sarcoidosis Biobanking per the University of Miami Institutional Review Board approval no 20150612. A written informed consent was received from participants prior to inclusion in the study. The animal study was reviewed and approved by Miami VA Animal Care and Use Committee (IACUC).

## Author contributions

MM conceptualized the study. MM and CZ designed the study. CZ, EMD, and RT conducted experiments. MM analyzed the data. AG performed RNA-Seq analysis. PB reviewed the lung tissues and performed tissue inflammation scores. AS and RC developed MIA602 and provided it for the experiments. MM, GH, and BJ interpreted the results. MM wrote the manuscript. All authors reviewed and edited the manuscript.

## Acknowledgments

Authors would like to acknowledge the Sylvester Comprehensive Cancer Center Flow Cytometry Shared Resource

Research reported in this publication was supported by the National Cancer Institute of the National Institutes of Health under Award Number P30CA240139. The content is solely the responsibility of the authors and does not necessarily represent the official views of the National Institutes of Health. This study was funded by VA Distinguished Scientist Award, VA (Grant No. 1 I01 BX004371), and VA Research Service.

## Supplemental Figure 1

**Figure S1.**
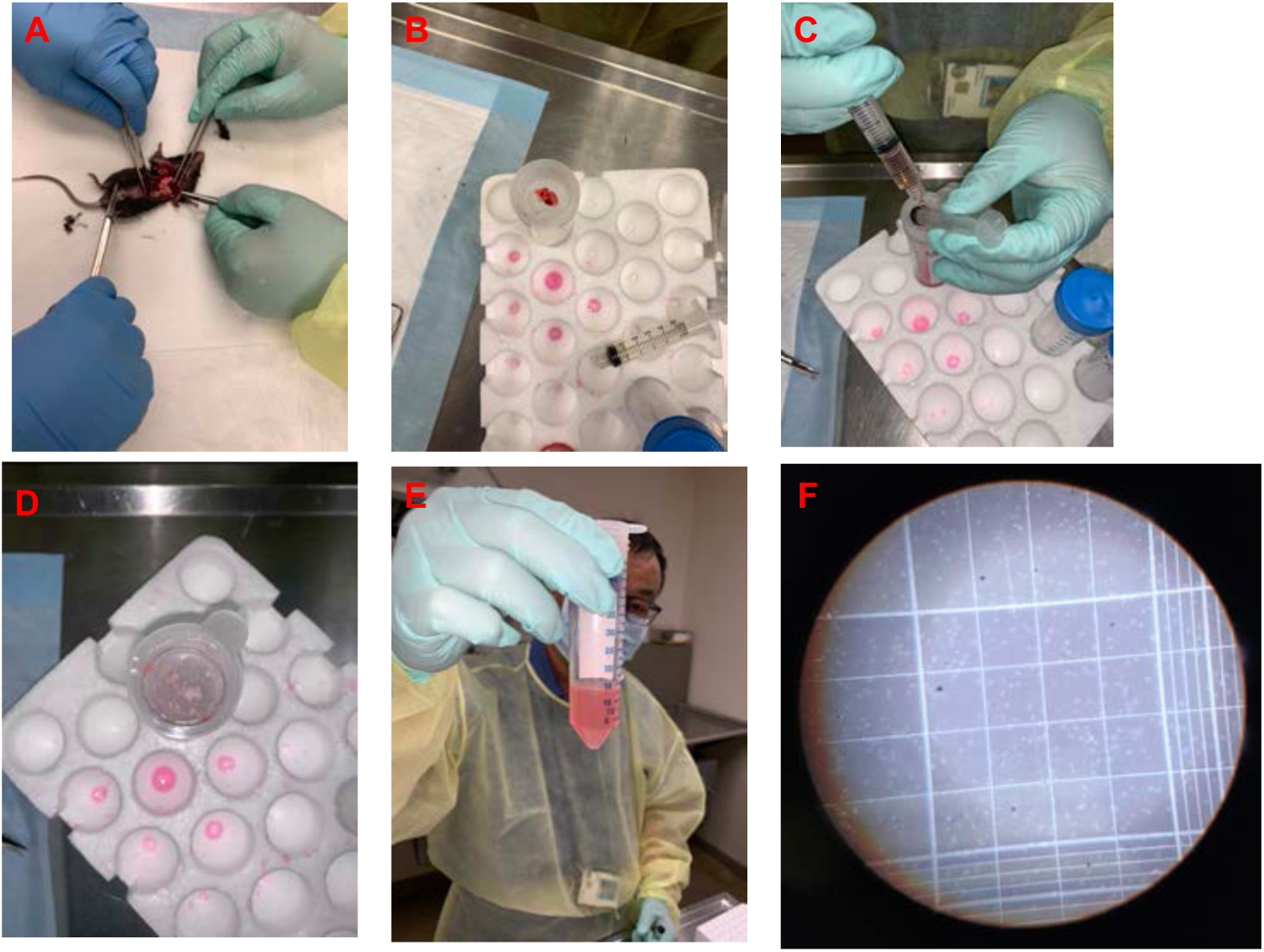
Shows the steps to develop single cells from mouse lung. A. remove lung tissue B. put tissue on sieve C. tissue was mashed with a rubber D. shows majority of cells already isolated and left elastin and collagen elements E. A cloudy fluid collected F. Shows isolated single cells under microscope.

